# Microbial Genome-Wide Association Studies: Lessons from Human GWAS

**DOI:** 10.1101/093211

**Authors:** Robert A. Power, Julian Parkhill, Tulio de Oliveira

## Abstract

The reduced costs of sequencing have led to the availability of whole genome sequences for a large number of microorganisms, enabling the application of microbial genome wide association studies (GWAS). Given the successes of human GWAS in understanding disease aetiology and identifying potential drug targets, microbial GWAS is likely to further advance our understanding of infectious diseases. By building on the success of GWAS, microbial GWAS have the potential to rapidly provide important insights into pressing global health problems, such as antibiotic resistance and disease transmission. In this review, we outline the methodologies of GWAS, the state of the field of microbial GWAS today, and how lessons from GWAS can direct the future of the field.

## Introduction

Over the last decade, genome-wide association studies (GWAS) have yielded remarkable advances in the understanding of complex traits and identified hundreds of genetic risk variants in humans (e.g. ^1-3^). GWAS normally analyse hundreds of thousands to millions of common genetic variants, usually single nucleotide polymorphisms (SNPs), and test for an association between each variant and a phenotype of interest (see ^4^). GWAS have confirmed the heritability of many human traits^5^, clarified their underlying genetic architecture^6^, and identified novel biological mechanisms and drug targets^7^. Of recent interest to infectious disease research are microbial GWAS, which identify risk variants on the genomes of microorganisms such as bacteria, viruses and protozoa. With increasingly cheap and high-throughput sequencing technologies, microorganism whole genome sequences (WGS) are now being generated on an unprecedented scale that rivals human data. Microbial GWAS provide a new opportunity to develop insights into the biological mechanisms underlying clinical outcomes, such as drug resistance and pathogenesis. As in human GWAS, insights from microbial GWAS may lead to the identification of molecular targets for drug and vaccine development. Furthermore, identifying genetic variants through microbial GWAS will allow researchers to track the evolution and spread of pathogenic strains in real time, and to synthesise microorganisms *in vitro* with desired clinical phenotypes.

Human GWAS provide an optimistic outlook for microbial GWAS. However, there are significant differences between microbial and human genomic studies that could hinder the success of microbial GWAS or require methodological adaptations. In this Review, we first outline specific features of GWAS methods and consider their application to microorganisms.

Second, we summarize the microbial GWAS that have been carries out to date, outlining their key findings, methods and challenges. Although these studies have focused mainly on pathogenic viruses, bacteria and protozoa, and thus dominate the focus of this Review, it is important to note that the same methods can be applied to non-pathogenic microorganisms. Finally, we discuss the lessons that have been learned from human GWAS and anticipate the future of microbial GWAS, particularly the opportunities provided by the ability to collect GWAS data from both the host and microorganis

## Data and methodology of GWAS

GWAS grew from the common disease common variant (CDCV) hypothesis^8^, which postulates that many high frequency but low effect variants contribute to disease risk. This explained how diseases can avoid selection, manifest in complex inheritance patterns, and be genetically and phenotypically heterogeneous. GWAS are implemented to identify the common variants that underpin the heritability observed for many phenotypes (see Box 1)^9^. These common variants are usually usually in the form of bi-allelic SNPs, where two nucleotides (A, C, G or T) exist at a locus with a frequency above 1% in the population. Each SNP is analysed, usually through linear or logistic regression, to determine whether one allele is significantly associated with the phenotype. Effects are reported as either beta for quantitative traits or odds ratio for case-control studies. Typically, only the main effects of individual SNPs are calculated, as methods for the detection of epistatic interactions between SNPs and SNP-environment interactions are challenging owing to the additional burden of multiple testing^10,11^. The power of the GWAS approach came from genotyping chips that enable the rapid calling of hundreds of thousands of SNPs from across an individual’s genome. Due to the co-inheritance of segments of the genome over generations, correlations (known as linkage disequilibrium, LD) exist between genetic variants in close proximity. LD allows genotyping chips to ‘tag’ local genetic variation by including a single proximal SNP, and to impute additional SNPs that were not directly genotyped based on known correlations^12^.

There are a number of conceptual differences between human and microbial GWAS (see Table 1), one of the most important of which is the source of the genomic data. Unlike human GWAS, where data comes from SNP genotyping chips, almost all genomic data for microorganisms comes from sequencing. This has an impact on several aspects of GWAS, particularly SNP calling, as SNPs detected in microbial sequencing data will not only be biallelic, but also tri- and quad-allelic. This complicates variant calling, data storage and analysis. Matching loci to a reference genome is also of increased importance in microbial GWAS, to ensure that SNPs are called at the same location for each sample and for comparison across studies. Sequencing also affects the quality control (QC) steps that must be taken to filter SNPs and individual samples. Standard QC in human GWAS removes those SNPs with low minor allele frequency (with a typical cut off of <1-5%), high missingness (>1-5%), or those that are out of Hardy-Weinberg Equilibrium (p<E-5 or-6). QC on individual samples in a human study also removes those samples with high missingness (>1-5%) or that are outliers in genome-wide homozygosity. Owing to the large number of SNPs compared to the number of samples in a study, QC is carried out to preferentially exclude SNPs. With the exception of Hardy-Weinberg Equilibrium, these same QC metrics will remain important for microbial GWAS. However, QC thresholds need to be established for additional metrics that capture the quality of sequencing, such as sequencing depth and Phred scores.

**Table 1:** Summary of the conceptual and analytic steps of human GWAS and microbial GWAS.

|  | **Human** | **Microorganism** |
| --- | --- | --- |
| Estimation of heritability | Twin studies<br>Adoption studies<br>GREML analyses | Within transmission pair correlations<br>Phylogenetic studies |
| Main source of GWAS data | SNP genotyping chips | Whole genome sequencing (WGS) |
| Common study designs | Case-control<br>Quantitative traits | Binary and quantitative traits<br>Longitudinal within individual sampling |
| Quality control steps | Individual samples missingness<br>SNP missingness<br>Hardy Weinberg Equilibrium<br>Minor allele frequency | Sequencing depth<br>Poor assemblage<br>Minor allele frequency |
| Reference genomes for imputation and linkage disequilibrium | International HapMap Project<br>1000 Genomes Project | RefSeq genomes<br>LD can be determined directly from sample |
| Confounding | Ethnic ancestry<br>Population stratification<br>Cryptic relatedness | Subtypes or Genotypes<br>Selection sweeps<br>Recombination and horizontal gene transfer<br>Clonal expansion |
| Significance Threshold | $p=5E-8$ | Differs by species<br>Currently no field-wide definition |
| Replication | Required for publishing novel associations | Not yet universally performed<br>Possibility of *in vitro* validation |

SNP, single nucleotide polymorphism; LD, linkage disequilibrium

## Adapting GWAS to microbial variants

As mentioned above, human GWAS typically focus on the effects of individual SNPs. However, focusing on the additive effects of SNPs alone will not always be possible in microbial GWAS. For instance, in bacteria recombination can introduce novel genes. This means the causative genetic difference may be the presence or absence of an entire gene or set of genes. Microbial GWAS need to test this variation in gene presence alongside SNPs. Here, lessons may come from the analysis of copy number variants (CNVs) in human GWAS. CNVs are large duplications or deletions of sections of the genome. CNV analyses test for associations between a phenotype and both specific CNVs and (as specific CNVs are often rare) an individual’s CNV burden. An individual’s CNV burden is the percentage of their entire genome, or a region of it, that is covered by CNVs^13^. Similarly, analyses of human sequence data often test for associations with the burden of rare variants^14^. The contribution of variants to that burden can be weighted by their predicted functional impact. Using quantitative burdens that combine the effects of multiple genetic variants into a single variable might provide statistical methods for analysing gene presence or absence and rare variants in microbial GWAS.

Another approach to handling gene presence in microbial GWAS is defining and analysing k-mers, a specific sequence of bases^15^. The benefit of k-mers is that they capture common variation and gene presence simultaneously. Analysis of k-mers may also be useful owing to the larger proportion of coding sequence found in many microorganisms, compared with humans where only a small fraction of DNA is exonic. This is because k-mers can capture multiple allele differences that code for different amino acids, and thus reflect changes closer to the biological mechanism that underlies the phenotype of interest.

It is worth noting that the majority of human GWAS have focused on additive effects of variants. This is where each additional copy of an allele carried by a diploid organism increases the likelihood of a phenotype in a linear manner. However, owing to within host evolution and the possibility of superinfections some microorganisms will exhibit within host genetic diversity. Within host diversity will lead to non-discrete SNP calling, where the frequency of an allele reflects its frequency on microbial sequences within the host, rather than an allele’s presence or absence. While testing for a linear association between allele frequency and phenotype makes pragmatic sense, the possibility of non-linear effects exists. Further, within host diversity results in alleles from different lineages having unique LD patterns. This will be relevant to the analysis of epistatic interactions, as alleles within the same host may have different genetic backgrounds.

Lastly, microbial GWAS are also likely to observe lineage effects. Here, entire lineages, such as viral subtypes, might differ in phenotype. In this instance, the lineage or subtype of the microorganism might be the genetic unit of interest, either alone or in addition to individual SNPs or k-mers. Disentangling the effects of a single variant from those related to lineage is potentially challenging, but has been shown to increase the power of microbial GWAS when implemented successfully^16^.

## Confounding factors in microbial GWAS

The main challenge associated with GWAS is the risk of identifying seemingly causal variants that are in fact false positives^17^. This is due to two main causes: population structure and multiple testing (see below). Recruitment of samples from within a genetically diverse population can lead to subtle confounding from population structure, e.g. because of an excess of cases from one ethnic group. In such instances, GWAS would identify predictive SNPs informative only of ancestry, not the biology of the disease. To avoid this problem, human GWAS often restrict recruitment to ethnically homogenous groups. Even within relatively homogenous populations, population structure exists. These subtler influences of population stratification are corrected through principal component analysis. This generates covariates that capture SNP correlations across the genome, and can be carried out using software such as EIGENSTRAT^18^. Principal components can capture subtle ancestry differences with high accuracy and identify samples that represent population outliers^19^. Although principal components will be key to removing confounding due to population structure in microbial GWAS, two additional confounders exist that may require additional methods than those used in human GWAS.

The first of these is homologous recombination, which occurs in bacteria and viruses through the replacement of short sequence blocks, rather than through multiple cross-overs along the whole chromosome. This means that long-range LD is broken down differently in microbial genomes, leaving variants in long range LD with each other even when short range LD within a region is reduced^20^. This long-range LD will make identification of the causal variant problematic^21^. Methods designed for analysing historically ethnically mixed, or “admixed”, human populations may be useful here, because they make use of recombination patterns to identify associated loci^22^.

The second source of confounding is that microbial population structure can represent selection on the phenotype of interest, e.g. antibiotic resistance. Given the differences in frequency of recombination and selection across microorganisms, the consequent population structures are likely to range from purely clonal to nearly panmictic. In addition, the rapid spread of successful epidemic lineages may temporally reduce their recombination with the rest of the species. In microorganisms where there has been strong selection, it may be appropriate to use repeated samples from within a single host over time, such as comparing pre- and post-treatment sequences. However, this approach will not work for longitudinal phenotypes, such as the time taken to develop disease symptoms, or in microorganisms with low rates of evolution. In these studies, methods that use mixed models to account for relatedness^15^ or lineage effects^16^, or to identify signals of selection across the genome based on phylogenetic structure^23^ may have more traction than typical GWAS regression methods.

## Multiple testing and replication

Aside from confounding, the other major source of false positives is the multiple testing that is intrinsic to GWAS. The standard cut-off for an association to be considered statistically significant is p=0.05, which represents a 5% probability of random occurrence. However, testing hundreds of thousands of SNPs leads to tens of thousands of SNPs being significant at p<0.05 by chance alone. To account for the number of tests, a SNP must pass the genome-wide significance cut-off in order to be considered significant (see Box 2). This is usually p<5E-8 in humans^24^, which is approximately equal to the Bonferroni correction (a multiple testing correction) for the number of SNPs analysed in early GWAS. However, it continues to be used in more densely genotyped and imputed studies today. Additional SNPs included in GWAS through deeper genotyping or imputation are in high LD with those already known, and so the correlations between SNPs reduces the number of independent tests performed. Thus, understanding the level of LD between SNPs is important for calculating the correct threshold for genome-wide significance. Even with strict cut-offs for genome-wide significance, determining if an association represents a false positive remains problematic.

As a result, replication in an independent cohort is the gold standard for reporting an association in GWAS^25^. This is both to avoid false positives and to accurately estimate the effect size of the SNP. Normally, GWAS have reduced power to detect variants of small effect and consequently there is a bias towards identifying novel SNPs that have an over-estimated effect size (sometimes called ‘Winner’s Curse’)^26^. As no bias for discovery exists during replication, the effect size in the replication cohort will more accurately reflect the true effect. Generally, replication does not require a SNP’s an association to reach genome-wide significance in the replication cohort, but to pass a p-value threshold based on for the number of SNPs brought forward for replication. Further, meta-analysis of the p-values of a SNP in both the discovery and replication cohorts should surpass genome-wide significance in order for a SNP to be considered a true positive.

However, microbial GWAS may be less reliant on replication than human GWAS given that suspected causal variants can be validated *in vitro.* This ability to generate carriers of identified variants and test their effect in the laboratory reduces many of the concerns of false positives that are typically associated with human GWAS. It also provides model organisms that can be used to gain a better understanding of the variant’s function. One important area of research is to account is the development of methods to identify and correct for epistasis. Epistasis can take the form of specific interactions between two SNPs or the effect of a SNP being conditional upon a broader genetic background. Disentangling epistatic effects will be key to generating viable *in vitro* models of microbial GWAS findings and establishing causality.

## Power, polygenicity and heritability

As well as providing methodological insights, the history of GWAS predicts a clear trajectory for how progress in microbial GWAS is likely to unfold. Initial human GWAS identified only a small number of SNPs, each explaining only a tiny fraction of variation. The disparity between expected heritability from twin studies and the heritability explained by genome-wide significant associations became known as the “missing heritability”^27^. Missing heritability initially cast doubt on the GWAS approach. Yet, as the first waves of studies were pooled into meta-analyses^28^ and the second waves of GWAS were analysed, more and more associations were reported, increasing the amount of heritability explained^29^. It became clear that the stringent cut-off for statistical significance resulted in a requirement for larger sample sizes than had been expected in order to achieve sufficient power to identify SNPs. Once sufficient power was reached, the relationship between the sample size and number of SNPs identified became relatively linear. However, despite this, there was often an inverse relationship between the frequency of identified SNPs and their effect size, meaning that each SNP explained only a small fraction of variation^29^.

The problem of missing heritability persisted, leading to a move away from single SNP analyses and towards polygenic methods^30^ (see Figure 1). One of the first polygenic methods was the use of polygenic risk scores (PRSs)^31^. PRSs are based on the assumption that many SNPs with small effect sizes will fail the stringent cut-off used for genome-wide significance, yet, together their cumulative effect could explain a large amount of the variance in risk. The construction of a PRS requires both a discovery and a replication cohort. In the discovery cohort, a GWAS is carried out, defining the ‘risk’ allele and effect size of each SNP regardless of whether the p-value is significant. In the replication cohort, the number of ‘risk’ alleles that an individual sample carries is summed into a score (the PRS), with each allele weighted by its effect size. The variation in case-control status predicted by the PRS is then calculated. Several PRSs are often defined using different p-value thresholds for the inclusion of SNPs from the discovery GWAS, e.g. four scores using SNPs with p<0.001, p<0.05, p<0.2, and p<0.5. As more SNPs are included, there is a greater likelihood that all SNPs of true effect will be included. However, including more SNPs also increases the number of SNPs with no true effect, and thus adds noise, which causes the amount of variance explained to plateau. PRSs ultimately provide a more powerful predictive tool than the results of single SNPs. As such, PRSs may be key to rapidly translating the results from microbial GWAS to prediction in the clinic, even before the roles of causal risk variants are understood.

**Figure 1:**
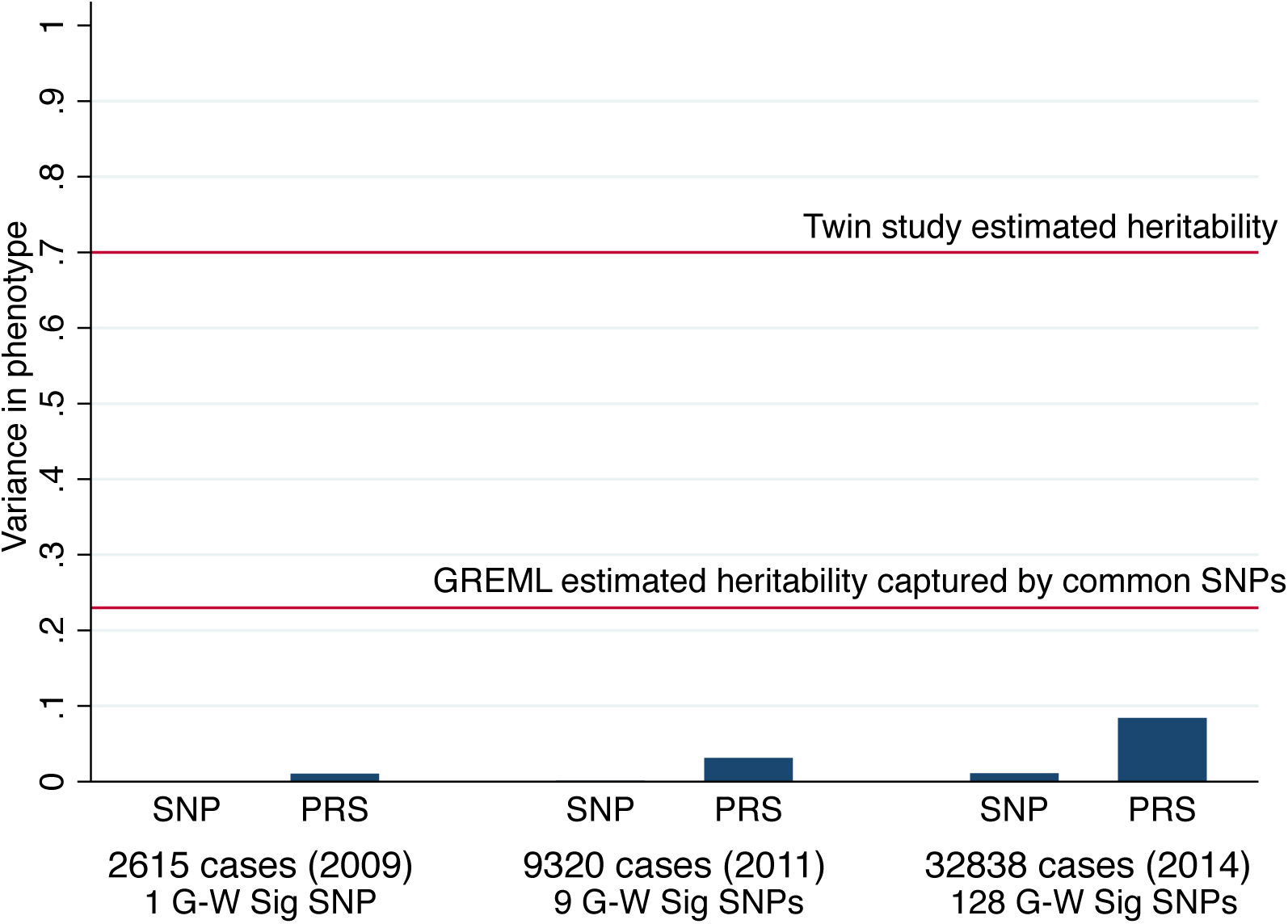
Phenotype prediction as GWAS sample sizes increase. Variance in a phenotype (schizophrenia^3^) explained in successive waves of GWAS studies by the genome-wide significant SNPs (SNP) and polygenic risk scores from all SNPs with p<0.05 (PRS). The number of genome wide significant SNPs (G-W Sig SNPs) identified also increases exponentially with sample size and at every stage PRSs provide substantially better prediction than the use of significant SNPs alone. However, the challenge of ‘missing heritability’ continues even within relatively large GWAS, with the variance explained still below the heritability estimates derived from GREML and twin studies.

An alternate polygenic method is genomic-relatedness-matrix residual maximum likelihood analysis (GREML), which was often referred to in the early literature by the software name GCTA^5^. GREML estimates the proportion of variance that is captured by all SNPs and calculates the heritability of the phenotype. This is done by calculating how genetically similar each possible combination of two samples is (i.e. their genetic relatedness). Relatedness refers to how much of the genome is shared between two samples (i.e. have the same genotypes). The heritability is then calculated as the proportion of phenotypic similarity between samples that can be explained by their relatedness. It is important to note that GREML does not estimate the true heritability of a phenotype, only the heritability that is captured by the included SNPs. Unlike PRS, GREML does not provide a means of predicting risk. However, it does act as a benchmark for the maximum amount of risk detectable in an infinitely powered GWAS. For example, in humans, GREML estimated that common SNPs account for between a third and a half of the heritability estimated from twin studies^30^(Figure 1). While PRS and GREML have not been widely used in microorganisms, they will be key to understanding whether current microbial GWAS are underpowered and if novel variants will be identified with larger sample sizes.

A crucial aspect of polygenic methods is their ability to identify what drives the heritability of a phenotype. First, polygenic methods can be used to test if heritability is disproportionately driven by specific genomic regions, by rare or common variants, or by variants within particular biological pathways. Second, polygenic methods can measure the heritability of specific subtypes of the phenotype. Identifying if a phenotypic subtype has higher heritability identifies those individuals for which the microbial genome is most relevant. Furthermore, polygenic methods are able to identify a genetic correlation between two phenotypes, even when data is available on only one phenotype in each sample^32^. Thus they can determine if two distinct phenotypes have overlapping aetiology, or if two subtypes of a phenotype are genetically distinct. Polygenic analyses have supported the generalist genes hypothesis, according to which genetic effects are highly pleiotropic^33^. Overall, human GWAS predict that for traits under moderate selection the genetic architecture will consist of many small effect and pleiotropic variants, which are spread fairly evenly across allele frequencies and genomic regions.

## Progress in microbial GWAS

Given the clear trajectory of human GWAS from underpowered studies to more advanced methods that explain a significant proportion of risk, it makes sense to ask whether microbial GWAS will advance in the same manner. Despite the complexities previously mentioned, a growing number of microbial GWAS have recently been published (Table 2). With the exception of HIV and *Plasmodium falciparum,* these publications have largely focused on bacteria and almost exclusively focused on pathogens within human hosts. The majority of genomic data has come from WGS, although genotyping chips for *P. falciparum* have existed for several years^34,35^. Owing to the much shorter genomes of microorganisms, the number of variants analysed in microbial GWAS has been in the tens of thousands, orders of magnitude smaller than in human GWAS. Sample sizes have also been significantly smaller. The smallest microbial GWAS to date was a study of 75 *Staphylococcus aureus* strains^36^ and the largest, a study of 3,701 *Streptococcus pneumoniae* isolates^37^. The majority of studies had sample sizes below 500 (see Table 2). However, this promises to change as large multicountry consortia, such as MalariaGEN^38^ and PANGEA_HIV^39^, generate whole genome sequences on a much larger scale.

**Table 2:** Examples of microbial GWAS to date.

| Organism | Genome length | Recombination rate | Within host diversity | Reference | Sample Size | Phenotype | N SNPs | Significant SNPs | Software |
| --- | --- | --- | --- | --- | --- | --- | --- | --- | --- |
| *Campylobacter jejuni* | 1.6 Mb | High | High | <sup>42</sup> | 192 | Host preference | - | 7,307 kmers in 7 genes | bespoke |
| *Mycobacterium tuberculosis* | 4 Mb | Low | Low | <sup>23</sup> | 123 | Drug resistance | 24,711 | 50 | PhyC |
|  |  |  |  | <sup>47</sup> | 123 | Drug resistance | 24,711 | 133 | PLINK |
|  |  |  |  | <sup>40</sup> | 498 | Drug resistance | 11,704 | 12 | PhyC |
| *Staphylococcus aureus* | 2.9 Mb | Low | Low | <sup>36</sup> | 75 | Drug resistance | 55,977 | 1 | ROADTRIPS |
|  |  |  |  | <sup>44</sup> | 90 | Virulence | 3,060 | 121 | PLINK |
| *Streptococcus pneumoniae* | 2.2 Mb | High | Low | <sup>37</sup> | 3,701 | Drug resistance | 392,524 | 301 | PLINK |
| *Plasmodium falciparum* | 22.9 Mb | High | Low | <sup>41</sup> | 1,063 | Drug resistance | 18,322 | 9 | FaST-LMM |
| HIV | 9,000 bp | High | High | <sup>45</sup> | 343 | Drug resistance | 5,100 | 8 | PLINK |
|  |  |  |  | <sup>43</sup> | 1071 | Viral load | 3,125 | 0 | PLINK |

Despite the current small sample sizes, microbial GWAS have already been successful in identifying causal variants. This is in part due to the studies focusing on phenotypes under strong selection, the majority of which were studies on drug resistance. For example, microbial GWAS of *Mycobacterium tuberculosis*^40^, *S. aureus*^36^, *S. pneumoniae*^37^, *P. falciparum*^41^, and HIV have all successfully identified novel drug resistance variants that often explained almost all of the phenotypic variation. Even with phenotypes under strong selection, there has been evidence of high polygenicity within microorganisms. For example, the study of drug resistance in 3,701 *S. pneumoniae* sequences identified 301 significant SNPs, with a median odds ratio of 11^37^. Given the large effect sizes, it is not surprising that many of the drug resistance variants identified through microbial GWAS were previously known. Though this diminishes the novelty of the findings, it also strengthens confidence in the ability of microbial GWAS to correctly identify causal variants. Another phenotype under strong selection is host specificity. Microbial GWAS of host specificity have yielded significant results, for *Campylobacter jejuni*^42^ and HIV^43^. However, within the same study of HIV host-specificity, the authors found no associations between viral variants and infectiousness. The most successful study of virulence was of 90 *S. aureus* samples^44^. The authors identified 121 SNPs at genome-wide significance. Functional follow up of a subset of SNPs showed that 4 out of 13 affected toxicity *in vivo,* suggesting that a proportion of associations identified were truly causal.

Most microbial GWAS to date have focused on the analysis of traits that are under strong selection, yet these studies have shown remarkable diversity in their analytic approaches (Figure 2). Two analyses of HIV sequences have been performed ^43,45^, both using the GWAS software PLINK^46^. Based on fixed effect models these studies suggested that the virus shows low levels of population stratification. However, analyses of *M. tuberculosis* highlighted that while PLINK could identify many drug resistance variants, it also led to false positives due to confounding from population structure^47^.To address this limitation, the authors developed the software PhyC^23^, a tool that uses phylogenetic trees to identify SNPs under recent convergent evolution. This approach identified many of the same drug resistance variants as PLINK, yet reduced the level of confounding from population structure. Other studies have included phylogenetic structure as a random effect in mixed models, using software such as ROADTRIPS^48^ and FaST-LMM^49^. These mixed models have successfully reduced the effect of population structure in a number of microorganisms^36,41^. One of the limitations of these software is that they are designed for human genomic data and cannot handle features such as within host microbial diversity. A recent study developed a bespoke approach to microbial GWAS in the analysis of *C. jejuni*^42^. The authors generated multi-allelic k-mers, rather than SNPs, and tested these for an association with host preference. This is the only study, so far, to combine analysis of SNPs and gene presence/absence, a key genomic feature of bacteria.

**Figure 2:**
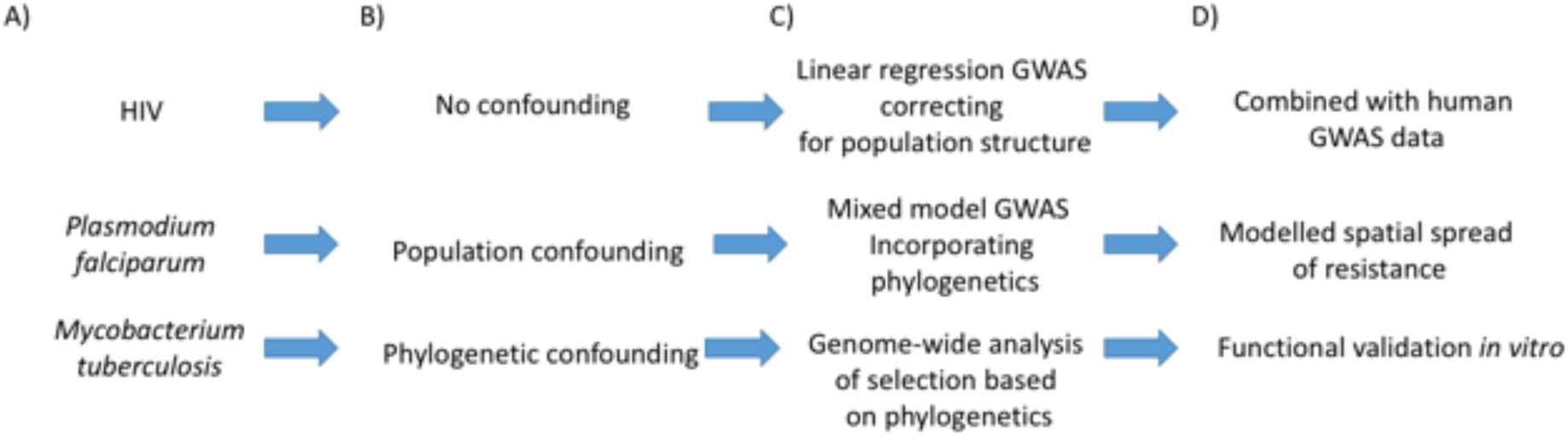
Potential models for microbial GWAS. Examples of three microbial GWAS approaches to date^40,41,43^. A) This panel shows the organism analysed in each study: HIV, a retrovirus that causes AIDS; *Plasmodium falciparum,* a parasitic protozoa that is the cause of malaria; and *Mycobacteria tuberculosis,* a bacteria that causes tuberculosis. B) This panel highlights the the form of geographic, population or phylogenetic confounding observed in each organism, which hinders the ability to differentiate SNPs of true effect from systematic false positives. For HIV only minimal population structure was observed, while for *P. falciparum* greater population differences existed. *M. tuberculosis* showed the highest level of confounding, with the different phenotypes (represented by the red/white nodes of the phylogenetic tree) clustering largely within the same lineages. C) Given the different population and phylogenetic structures of the three organisms, three different approaches were used to perform the microbial GWAS. The lack of confounding in HIV allowed for the application of typical human GWAS fixed effect models. The more substantial population structure in *P. falciparum* was accounted for by including phylogenetic relatedness as a random effect in a mixed model. Lastly, the clear phylogenetic structure of *M. tuberculosis* was used to perform genome-wide analysis of convergent selection. D) This panel highlights how the results of each microbial GWAS were taken forward to better understand the microorganism. For HIV, the viral genomic data was combined with human GWAS data to perform a genome-to-genome analysis of HIV viral load. For *P. falciparum,* the information on drug resistance variants was combined with geographic data to highlight the spread of resistance variants through South East Asia. Lastly, for *M. tuberculosis,* the identified drug resistance variants (delta-ald and delta-ald-comp) were functionally validated by showing carriers had improved growth comparable to other resistance strains (BCG), outperfoming the wild type (WT).

Overall, it is clear that while microbial GWAS are yielding important insights into infectious disease, the field has yet to settle on a consistent analytic approach and current methods are not yet ideally suited to microbial genomes. More refined analytical methods will become particularly important as the focus of microbial GWAS expands beyond drug resistance and towards phenotypes where variants have subtler polygenic effects.

## Remaining lessons

As microbial GWAS become more widespread, there are still several lessons that can be learned from human GWAS. Perhaps the most crucial lesson revolves around the generation of sufficient sample sizes to identify variants of small effect. This requires a collaborative approach. Samples must often be pooled from across the world in order to create well powered discovery and replication cohorts. Of particular note is the mega-analytic approach that pools raw genotype data from all sites into a central repository, which is used for standardised QC and to increase power^50^. There is good reason for optimism here, as international microbial research consortiums already exist.

One area that has not yet been explored in microbial GWAS is the trade-off between sample size and heterogeneity. As more complex phenotypes are analysed, heterogeneity will reduce power to detect the causal variants. With finite resources and time, typically choose between focusing on collecting detailed clinical data on a smaller number of more homogenous samples, or recruiting large numbers of samples with minimal screening. In human GWAS both approaches have been shown to be effective. Firstly, power can be improved by restricting to “super controls”^51^, e.g. using controls on the opposite extreme of the phenotype of interest, or focusing on a subset of samples with a phenotype believed to be more homogenous or heritable^52,53^. Secondly, ‘minimal phenotyping’ can be used to maximise sample size, such as assuming all those with records of treatment are ill^54^. Widely collected proxy phenotypes, such as education level as a proxy for cognitive ability, have been successfully used to maximise sample sizes for more complex traits^55^. Aetiologically similar phenotypes can also be jointly analysed to maximise sample size^2,56^. Overall, a sensible first step appears to be to increase sample sizes as much as possible. This can then be followed by secondary analyses of more homogenous phenotypic subtypes where data is available.

Finally, many advances in human GWAS were made possible by free and open software applications (such as GCTA^5^ and PLINK^46^) that could handle a variety of data formats and perform multiple analyses (Table 3). These software applications were generally very user friendly, with detailed documentation. To date, microbial GWAS have been performed using a range of software with different analytic approaches (see Table 3). Although GWAS software that can handle large genomic datasets already exists, they are not ideally suited to the non-diploid multi-allelic nature of some microbial genomes. Nor can they perform longitudinal within-individual sequence comparisons that might be desired. In particular, GWAS methods will need to be adapted to deal with within-host microbial diversity and recombination. Further, the successful polygenic methods for estimating the heritability and co-heritability of phenotypes from GWAS data have yet to be evaluated in microbial GWAS. As can be seen from GCTA^5^, a single piece of software with a topical application has driven a large number of high profile advances in human genomics. The development of free and open software applications that can accurately and conveniently analyse a wide range of microbial WGS data to detect single SNP and polygenic effects is, therefore, a top priority of the field.

**Table 3:** Features of software applications used in microbial GWAS to date.

| **Software** | **Reference** | **Analysis** | **Population structure adjustment** |
| --- | --- | --- | --- |
| PLINK | 46 | Linear/logistic regression of allele count at SNP | Ancestry informative principal components and other covariate inclusion |
| PhyC | 23 | Identifies SNPs undergoing recent convergent evolution | Based on phylogeny, so inherent |
| ROADTRIPS | 48 | Association analysis of SNP effect, allowing random variables to account for sample relatedness | Corrects for provided or derived relatedness between samples |
| FaST-LMM | 49 | Association analysis of SNP effect, allowing random variables to account for sample relatedness | Derives relatedness matrix and corrects as random effect. Principal components can be included as covariates. |
| SEER | 15 | Linear/logistic regression using k-mers, simultaneously testing SNPs and gene presence/absence | Identifies relatedness from data using multidimensional scaling and generates covariates for regression |

### Future directions: integrating the host

Arguably, the most exciting application of microbial GWAS is to integrate it with human genomic data. Human GWAS of infectious disease have been performed for over a dozen pathogens (reviewed here^57^). This review will end by highlighting the potential for combining these findings with those of microbial GWAS. These genome-to-genome analyses can give important insights into whether the effects of microbial variants are universal or dependent on a specific host genetic background. Such statistical host-microbial interactions would help identify which host proteins the microorganism was interacting with on a molecular level. Further, interactions that prevent infection or disease progression would represent potential drug or vaccine targets.

To date, we are aware of only one comprehensive genome-to-genome analysis. The microbial GWAS of HIV set point viral load, previously mentioned, generated both HIV sequences and host GWAS data^43^. This study was able to identify many associations between viral genetic variants and those in the human genome, specifically within the major histocompatibility complex region. In a secondary analysis, the authors also highlighted the importance of host-pathogen correlations and how they might lead to overestimates of the combined host and pathogen heritabilities^58^. In this case, while both host and viral heritability of HIV set point viral load was observed, the two were shown to substantially overlap.

With cheaper genome sequencing methods, the ability of groups to generate both host and microbial data on the same individuals will only increase. However, just as microbial GWAS currently lack universal analytic software, so do genome-to-genome analyses. Such statistical tools will be needed in order for the field to flourish, particularly as the scale of data will make these analyses computationally intensive. A simpler method may be to condense multiple SNPs into a single variable, as seen in PRS^31^, and test for interactions on a genome-wide level. Regardless of method, the availability of host and microorganism GWAS data presents an opportunity to increase power to identify causal variants. Ideally, such data will be generated within large longitudinal studies, where genomic data can also be combined with epidemiological and clinical variables. Understanding the correlations between host demography, host heritability and microorganism heritability will give greater insight into the extent to which microbial genomes drive risk.

## Conclusions

As this review has shown, there is great promise in the field of microbial GWAS. However, it is clear that a number of analytical advances will be needed to handle the unique features of microbial genomics. Perhaps the issue of greatest importance will be the development of software applications that can handle the combined analysis of host and microorganism genomic data. With these tools, we will be better able to predict individual patient outcomes, track the evolution of global epidemics, and identify new drug and vaccine targets.

### Box 1: Heritability

The goal of GWAS is to identify the variants that determine heritable phenotypes. Heritability is the proportion of variation in the phenotype attributable to inherited genetic similarity. Knowing the heritability of a phenotype provides practical advantages to microbial GWAS. It provides an upper limit to the extent to which the phenotype can be predicted by identified variants. For some phenotypes the heritability may be obvious, such as antibiotic resistance being the result of drug resistance mutations^59^. For other phenotypes, such as HIV set point viral load, there has been debate regarding the extent to which viral genetic variants play a role^60^. Microbial heritability can be established in two ways. First, by looking at the correlation in phenotype across chains of transmissions. This determines the extent to which the same microbial variants lead to the same phenotypes across individuals. Second, by estimating the extent to which phylogenetic relatedness predicts similarity in phenotype. This determines the extent to which genetically similar microorganisms are phenotypically similar.

However, heritability estimates come with several caveats. First, there is a discrepancy between what is “genetic” and what is heritable. For example, a *de novo* genetic mutation would not be captured within heritability estimates nor would two identical changes on an amino-acid level that differed on a genetic level. Second, microbial heritability, host heritability and the environment explain the total variation in phenotype in a population. As a result, microbial heritability is relative to the amount of environmental and host variation. As the host and environment becomes more homogenous the microbial heritability increases, and vice versa. The heritability of a phenotype can change, or remain the same, independently of whether the mean value of the phenotype changes over time. Lastly, studies often estimate only additive genetic effects (known as narrow sense heritability), assuming no interaction between genes either at a single locus (dominance) or between loci (epistasis). However, uncovering epistatic interactions will be key to microbial GWAS in order to disentangle the effects of microbial variants from host background.

### Box 2: Visualising GWAS results

Two types of plot are used to visualise the results of genome wide association studies (GWAS). The first is the Manhattan plot, which plots each variant’s p-value against its position (see the figure; panel a). The x-axis represents the genomic location. The y-axis is the –log(p-value). The logarithmic scale is used so that the most significant SNPs stand out with higher values than the majority of non-significant SNPs. A reference line is used on the y-axis to reflect genome-wide significance, occasionally with a second line to represent a ‘suggestive significance’ threshold. Owing to the expectation of linkage disequilibrium (LD), a single highly significant SNP on its own is often interpreted as a genotyping error. Columns of significant SNPs in LD with the truly causal variant are seen in human studies, though this expectation is dependent on the LD of the organism.

The second is the quantile-quantile (QQ) plot, which compares the distribution of–log(p-values) observed in the study (y-axis) to the expected distribution under the null hypothesis (x-axis; see the figure; panel b). Departure of observed SNP p-values from the y=x reference line may reflect systematic inflation in the test statistics owing to population stratification. However, departure from this line is also expected for a truly polygenic trait, as many causal SNPs may not yet have reached genome-wide significance owing to lack of power. This will lead to an excess of low p-values across all SNPs. As a result, it is the point at which the observed –log(p-values) depart the y=x distribution that is important. Inflated –log(p-values) for all SNPs reflects population stratification, whereas polygenicity should lead to inflation for only those SNPs with high –log(p-values). The QQ plot is, therefore, a qualitative judgement rather than a quantitative one. However, a calculation of the lambda value (also known as the genomic inflation factor), which is derived by dividing the median value of the observed chi-squared statistic by the median expected chi-squared statistic (for p=0.5), gives a measure of the inflation in the sample. This should be 1 in the case of the null and is generally seen as inflation if above 1.05. The lambda value can be weighted by sample size to avoid polygenic inflation, as larger samples have the power to detect inflation due to many SNPs of small effect. In this case, a lambda value of 1000 is used to get an inflation estimate proportional to a GWAS that contained only 1000 samples.

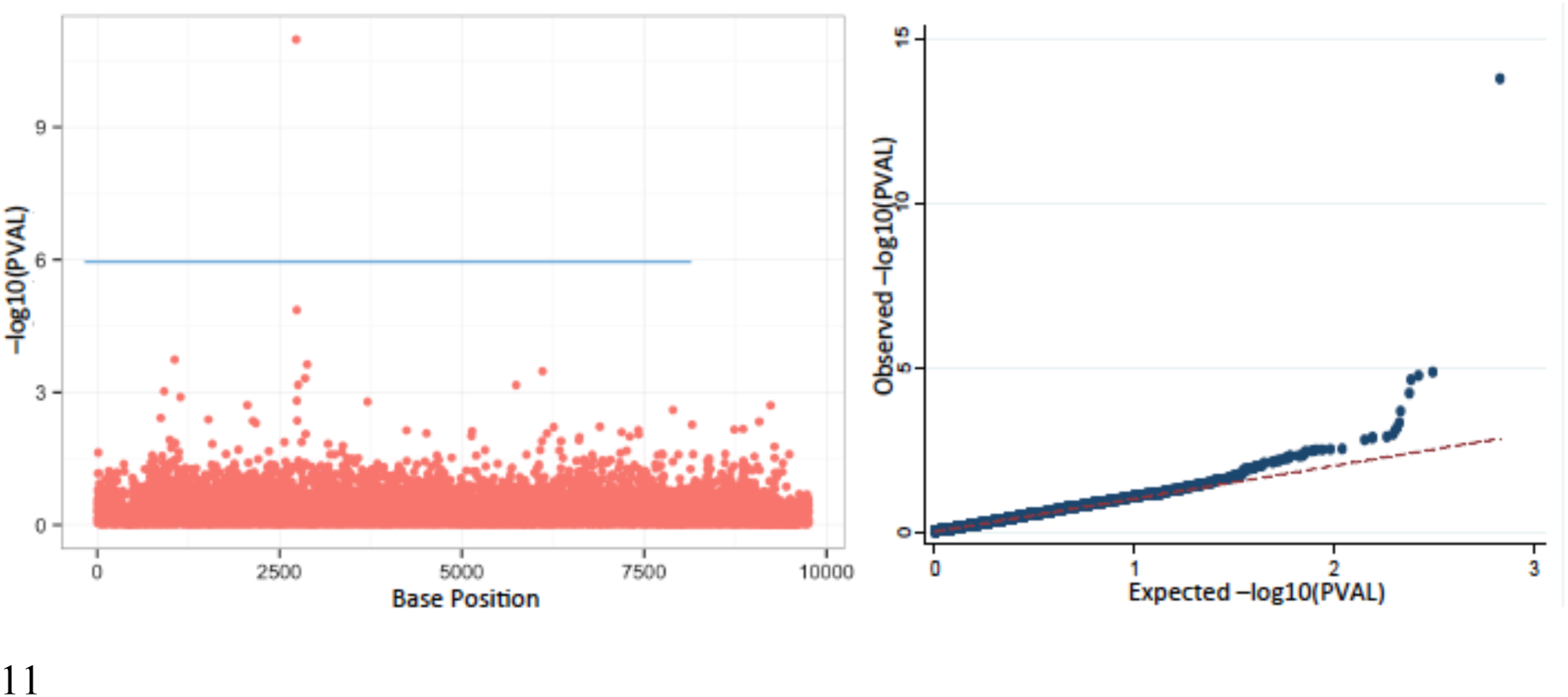

## Acknowledgements

Research supported by a South African MRC Flagship grant (MRC-RFA-UFSP-01-2013/UKZN HIVEPI) and Wellcome Trust grants (098051 and 201355/Z/16/Z.)

## Biographies

Dr Robert Power is a Sir Henry Wellcome postdoctoral fellow at the Africa Health Research Institute. His main research interest is the application of statistical genetic tools to both micro-organisms and humans to better understand disease aetiology. He received his PhD in statistical genetics from the Institute of Psychiatry, King’s College London.

Professor Julian Parkhill is Head of Pathogen Genomics at the Wellcome Trust Sanger Institute. He is interested in the use of high-throughput sequencing and phenotyping to study pathogen diversity and variation, particularly what they tell us about the evolution of pathogenicity and host interactions. He received his PhD in 1991 at the University of Bristol, was elected a Fellow of the Academy of Medical Sciences in 2009 and elected a Fellow of the Royal Society in 2014.

Professor Tulio de Oliveira is a professor at Nelson R Mandela School of Medicine at UKZN and a Royal Society Research Fellow at the Wellcome Trust Sanger Institute, U.K. He is recognized as an expert on HIV genetic data and bioinformatics software development. He received his PhD at the Neslon R Mandela School of Medicine in 2003 and was a Marie Curie Fellow at University of Oxford from 2004 to 2007. Tulio de Oliveira’s home page – www.bioafrica.net

## Key points

-GWAS have been highly successful in the analyses of human genomic data. The increased availability of microorganism whole genomes provides the opportunity for microbial GWAS.

-Initial microbial GWAS have had success identifying variants for traits under strong selection, such as drug resistance, in a range of bacteria, viruses, and protozoa.

-Several challenges to microbial GWAS exist that could hinder identifying variants under moderate selection. The primary challenge is the increased population stratification in microorganisms due to selection and complex recombination patterns.

-Novel software tailored to the needs of microbial GWAS would greatly expedite progress in the field. In particular, the application of polygenic methods has yet to be evaluated in microorganisms.

-An exciting future area of research is generation of host and microbial genomics data within the same samples. This will allow for genome-to-genome analyses to test for host-microbe interactions.

## Glossary

Beta: The standardized regression coefficient, derived from linear regressions in GWAS of continuous traits. It is reported as an estimate of the effect size of a SNP, and reflects the change in phenotype expected from carrying a copy of the reference allele of the SNP.
Clonal: Where reproduction produces genetically identical organisms, and so does not introduce novel variants or recombination.
Effect size: The proportion of variance in a phenotype predicted by a variant.
Epistatic interaction: Interactions between variants at different locations in the genome.
False Positive: A variant, or any predictor, that is identified as significantly associated with a phenotype but is not causal. In the case of GWAS this is usually due to confounding from population structure or insufficient quality control.
Genome-wide association study: A hypothesis-free method that tests hundreds of thousands of variants across the genome to identify alleles that are associated with a phenotype.
Genome-wide significance: The p-value cut-off for declaring a variant significantly associated with a phenotype, accounting for for the number of variants tested and the correlations between them.
Heritability: The proportion of phenotypic variance that is due to inherited genetic variation.
k-mers: A specific sequence of bases that, in microbial GWAS, can be used as the genetic variant tested for association with the phenotype.
Linkage disequilibrium (LD): Correlations between variants due to co-inheritance. LD is usually higher between variants that are closer together, and is broken down by recombination.
Main effect: The effect of a variant on the phenotype without accounting for any possible interactions with other variants or environmental factors.
Odds Ratio: The odds ratio, often abbreviated to OR, is the typical means of reporting the effect size of a SNP in a case-control (or other binary phenotype) GWAS. It is derived from a logistic regression, and represents the the odds of the phenotype when carrying the reference allele, compared to the odds of the phenotype in absence of the reference allele.
Panmictic: A population where all organisms are potential partners with each other.
Phenotype: A trait or disease that is the outcome of interest in an analysis of genetic variants.
Phred scores: A measure of the quality of sequencing at a given locus, specifically the confidence in the calling of alleles at that locus.
Pleiotropic: Pleiotropic variants are those that have an effect on multiple distinct phenotypes.
Polygenic methods: Statistical approaches that focus on the combined effects of many genetic variants rather than the effect of any individual variant.
Power: The probability that an analysis will reject the null hypothesis when the alternative hypothesis is true. Is influenced by numerous factors, such as the effect size and sample size.
Single nucleotide polymorphism (SNP): A base position where two alleles exist with a frequency of >1% in the population. Superinfection When an individual is infected with multiple strains of the same microorganism.

## Suggested highlighted References

4 –This review discusses in detail the methods, nuances and caveats of GWAS

15 –This methods paper presents a mixed model approach to microbial GWAS, including the analysis of k-mers

16 –This methods paper presents an approach to disentangling the effects of single SNPs and lineage effects within microbial GWAS

21 – The authors present an important review of the findings of bacteria GWAS to date.

29 –An important perspective on the lessons learnt from human GWAS and predictions of the future of the field.

30 – A useful review of a range of polygenic methods and their application.

23 – This microbial GWAS introduces the PhyC method that uses phylogenetic trees to perform a genome-wide scan of convergent evolution.

43 – An example of a genome-to-genome analysis with both host and microbial GWAS data.

